# ProGenFixer: an ultra-fast and accurate tool for correcting prokaryotic genome sequences using a mapping-free algorithm

**DOI:** 10.1101/2025.05.09.653025

**Authors:** Lifu Song, Mei Wang, Xiaoping Liao, Zhenkun Shi, Ruoyu Wang, Haoran Li, Lulu Liu, Botao He, Xiaomeng Ni, Jinshan Li, Hongwu Ma, Ping Zheng, Jibin Sun, Yanhe Ma

## Abstract

Although prokaryotic genomes are simpler, making variation analysis more straightforward, an efficient and user-friendly tool for their rapid sequence correction is lacking. We present ProGenFixer, an ultra-fast, mapping-free tool specifically designed to efficiently identify and correct errors in prokaryotic genomes using next-generation sequencing (NGS) data. ProGenFixer compares *k*-mer profiles between the reference genome and reads to pinpoint discrepancies, employs a local assembly-based algorithm to estimate corrected sequences, and automatically implements corrections. Benchmarking demonstrates ProGenFixer’s speed and accuracy advantages. It is over >5x faster than the swiftest existing tool tested while maintaining higher accuracy. Compared to slower, highly accurate tools, ProGenFixer shows comparable accuracy but is >17x faster and demonstrates enhanced performance on long indels (>10 bp). Implemented in C, ProGenFixer is a standalone, user-friendly tool offering a comprehensive solution for prokaryotic genome correction. The software is freely available at https://github.com/Scilence2022/ProGenFixer. Additionally, a companion web-based platform offering a ready-to-use interface can be accessed at https://progenfixer.biodesign.ac.cn.

## Introduction

The accuracy of genome sequences is becoming increasingly important to a wide array of biological research and applications, ranging from understanding microbial evolution and function to developing novel antimicrobial therapies and biotechnological processes [1–3]. Next-generation sequencing (NGS) technologies have revolutionized the field of genomics, enabling the rapid and cost-effective sequencing of entire genomes. However, the massive genome sequences generated by these technologies are inherently prone to errors arising from various sources during library preparation, sequencing, and data processing. Additionally, rapidly growing bacterial cells are prone to replication errors, which can further introduce more and more mutations in genome sequences. Since errors within coding regions, such as insertions and deletions (indels) which can shift reading frames, and substitutions which can alter key residues, can significantly impact protein function, correcting these errors in the genome sequence is critical to downstream phenotypic analyses. Therefore, genome sequence correction, also referred to as genome polishing, is essential for enhancing the precision and dependability of genomic data, thereby strengthening the validity of research findings.

Numerous genome sequence correction tools have been developed, employing diverse algorithmic strategies [4–10]. These tools can broadly be categorized into mapping-based and mapping-free methods. Mapping-based methods typically rely on aligning sequencing reads back to a reference genome sequence to identify discrepancies and correct errors. Recent advancements have led to the development of fast read aligners that utilize hash functions, advanced indices, and seeding techniques to accelerate the alignment process. Despite these advantages, mapping-based genome correction can be computationally expensive and time-consuming. In contrast, mapping-free methods operate by comparing *k*-mer profiles (short subsequences of length *k*) rather than aligning full-length sequences. The concept of *k*-mers, which has broad applications such as sequence alignment, genome assembly, and DNA data storage [11,12], plays a central role in these methods. Mapping-free algorithms are generally computationally inexpensive and can be significantly faster than mapping-based methods. Mapping-free methods also have limitations. For instance, accurately correcting errors within repetitive regions of the genome poses a significant challenge for these methods, as the *k*-mers derived from such regions are often ambiguous and occur in high frequencies [13,14].

Prokaryotic genomes are typically simpler than eukaryotic ones (haploid, smaller and fewer repeats), making variation analysis more straightforward. However, most existing genome correction tools are designed to handle eukaryotic complexities like polyploidy and extensive repetitive regions, leading to unnecessary computational overhead when applied to prokaryotic genomes. Additionally, these tools often impose further burdens on users by requiring dependencies on external read mapping and sorting software, which exacerbates both computational costs and installation difficulties. Thus, there is a clear need for an efficient and user-friendly tool specifically tailored for the rapid correction of prokaryotic genome sequences.

To address this need, we present ProGenFixer, a mapping-free tool engineered for rapid and accurate prokaryotic genome correction. ProGenFixer rapidly compares *k*-mer profiling of NGS data with the ordered *k*-mer array of a reference genome to detect low-quality regions. It then estimates the altered sequence content regions by regions using a local assembly algorithm. Unlike previous studies that use a single coverage threshold to eliminate low-coverage *k*-mers, ProGenFixer employs differential thresholds when detecting low-quality regions and estimating altered sequence content, balancing sensitivity and precision. ProGenFixer was implemented in multi-threaded C for maximum efficiency. Our benchmarks against leading tools including Jasper [6], NextPolish [7], Pypolca [8], and BWA+GATK [15,16], demonstrate ProGenFixer’s exceptional speed—more than five times faster— and its superior accuracy, especially in correcting long indels (>10 bp), establishing it as a valuable resource for prokaryotic genomics.

## Results and discussion

### Design and implementation of ProGenFixer

As illustrated in **Fig. 1A**, ProGenFixer operates using an iterative workflow consisting of five major steps, four of which are repeated in each iteration. Fundamentally, ProGenFixer refines a reference genome sequence by analyzing *k*-mer frequencies from the NGS reads and comparing them against *k*-mers derived from the reference sequence. This comparison pinpoints regions with discrepancies or low *k*-mers support, marking them as potential variation regions. To resolve these flagged areas, the tool constructs alternative sequences by navigating a *k*-mer graph weighted by *k*-mer counts, ultimately selecting the most probable correction. The resulting corrected genome sequence then serves as the input reference sequence for next iteration of correction, allowing for progressively enhanced accuracy. The five steps are detailed as follows:

**Fig. 1.**
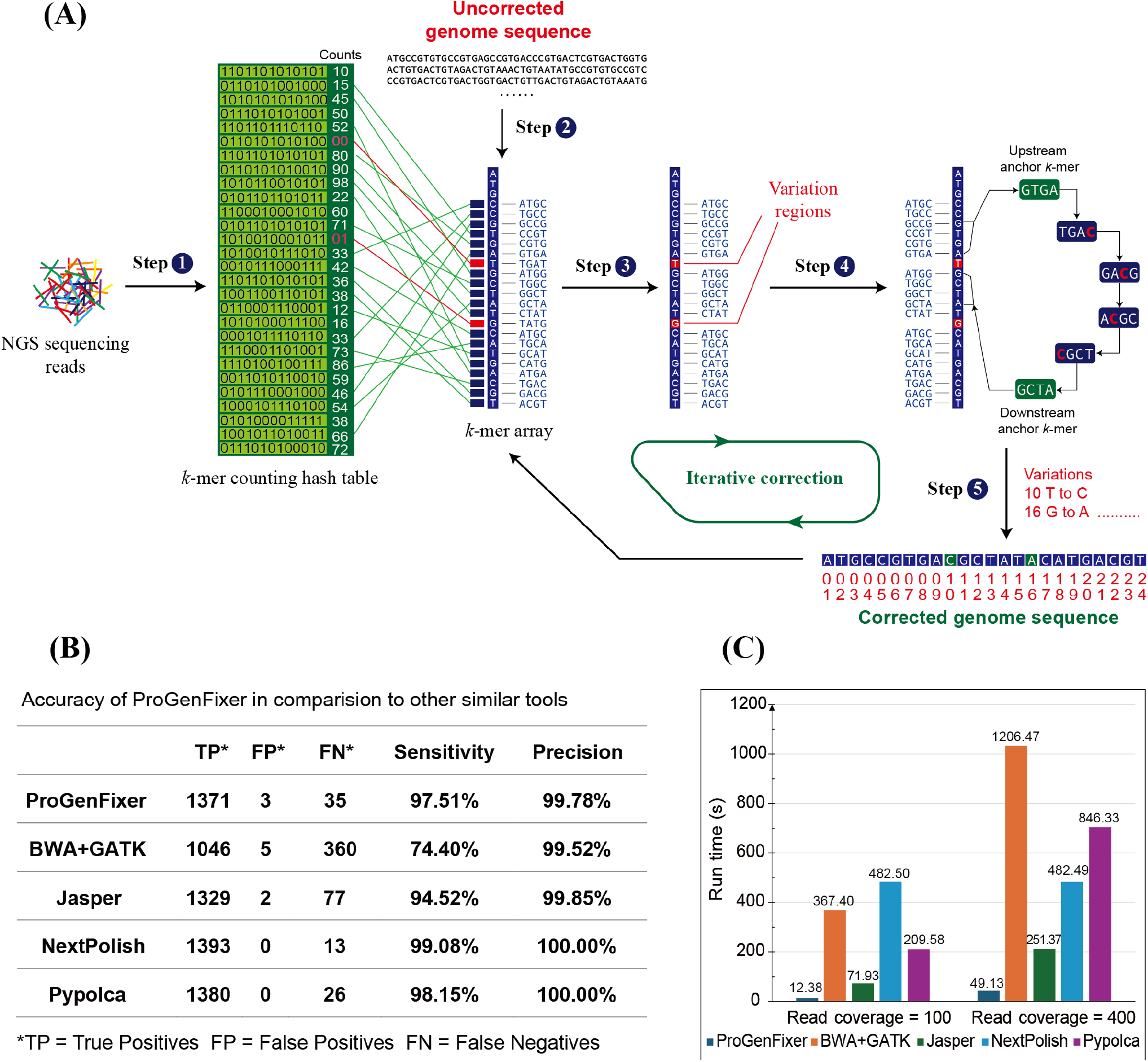
Workflow, accuracy and speed of ProGenFixer in comparison with existing genome correction tools. **(A)** Schematic overview of the ProGenFixer workflow. **Step 1:** Construct a *k*-mer counting hash table from NGS reads (without using a bloom filter). **Step 2:** Encode the reference genome sequence into an ordered *k*-mer array, mapping each *k*-mer to its genomic position. **Step 3:** Identify potential variation regions (highlighted in red) by checking the support for reference *k*-mers within the read-derived *k*-mer hash table. **Step 4:** Determine the corrected sequence content within variation regions using a local assembly-based approach. **Step 5:** Produce the final corrected genome sequence incorporating identified variations (illustrative example shows 10T→C and 16G→A changes). This corrected sequence can serve as the input reference for subsequent correction rounds (iterative correction). **(B)** Accuracy metrics (TP = True Positives, FP = False Positives, FN = False Negatives, Sensitivity, Precision) of ProGenFixer compared to other tools for correcting a mixture of substitutions and indels. **(C)** Speed comparison of ProGenFixer and other genome correction tools. The bar graph displays the run time (seconds) required to correct the *E. coli* MG1655 genome using various tools (ProGenFixer, BWA+GATK, Jasper, NextPolish, Pypolca) with simulated read coverages of 100X and 400X. Runtimes represent the average of three independent replicates; standard deviations are provided in **Table S2** (too small to visualize as error bars).

**Step 1**, ProGenFixer counts the *k*-mers of NGS reads using a simple and efficient hash table. This choice ensures that large datasets are processed swiftly while maintaining the accuracy of *k*-mer counts. Given the relatively small size of bacterial genomes, the advantages of the Bloom filter structure are limited, although it has been shown to be an effective strategy for reducing memory usage. As a result, the Bloom filter structure is not implemented in ProGenFixer. Additionally, the *k*- mer counting process is implemented using a multi-threaded architecture to enhance performance.

**Step 2**, the reference genome sequence is encoded into ordered arrays of *k*-mers, with each index correlating to the *k*-mer’s position within the genome sequence. This representation enables ProGenFixer to track the precise position of each *k*-mer in the reference genome, which is crucial for accurately detecting low quality regions with potential variations in the next step.

**Step 3**, ProGenFixer carefully examines each *k*-mer in the *k*-mer array derived from the reference genome sequence to assess its presence in the *k*-mer counter table generated from the NGS reads. *K*- mers that are either absent or have coverage values below a specified threshold in the counter table are flagged as potential variation regions, indicative of errors. To minimize amplification and sequencing errors, previous studies often implement a uniform *k*-mer coverage threshold to filter out low-coverage *k*-mers. In contrast, ProGenFixer utilizes differential thresholds during two consecutive processes: identifying variation regions and estimating the altered sequence contents (*i*.*e*., variation calling). A relatively low coverage threshold is used to identify variation regions, while a higher threshold is applied for estimating the altered sequence contents. This dual-threshold strategy ensures that the flagged variation regions are true sites of potential errors, minimizing the risk of introducing incorrect corrections.

**Step 4**, for each variation region identified with potential errors, ProGenFixer estimates alternative sequences by traversing the *k*-mer graph. The algorithm selects the most likely path based on the *k*- mer counter table. Specifically, ProGenFixer first merges variation regions that are within a distance not greater than the *k*-mer size. Next, the algorithm extends the region by including upstream and downstream high-quantity anchor *k*-mers (those present in the reference array and adjacent to the variation region), as illustrated in **Fig. 1A**. Two criteria must be satisfied for a *k-*mer to be selected as an anchor *k*-mer: first, it must not be present multiple times in the reference genome sequence; second, the corresponding coverage in the *k*-mer counter table must be higher than a specified threshold for estimating the altered sequence contents. If the nearby *k*-mers do not meet the anchor criteria, ProGenFixer will continue extending the region outward from the variation site, searching for the nearest *k*-mer that satisfies the criteria. Finally, the alternative sequence is estimated by traversing the *k*-mer graph using the counter table.

**Step 5**, ProGenFixer corrects the reference genome sequence based on the variations estimated in the previous step. The corrected genome sequence then becomes the new reference for the next iteration of corrections, as needed.

### Accuracy and speed of ProGenFixer in comparison with existing tools

We evaluated ProGenFixer’s performance using simulated data based on the genomes of two industrially important strains: *Escherichia coli* MG1655 (LR881938.1) and *Corynebacterium glutamicum* ATCC 13032 (BA000036.3) [1,2]. Results were highly consistent between the two genomes; therefore, we present representative results for *E. coli* MG1655 here, with complete simulation results available in **Supplementary Fig. S1** and **Tables S1-S4**.

First, we assessed the performance of ProGenFixer in handling a mixture of substitutions and insertions/deletions (indels). Using *dwgsim* v0.1.15 [17], we simulated 150 bp paired-end NGS reads at 100X and 400X coverage. The simulation incorporated a mutation rate (-r) of 0.3‰ (parts per thousand), with 10% of mutations being indels and an indel extension probability of 0.3. To benchmark ProGenFixer, we compared its performance against established tools: BWA+GATK, Jasper, NextPolish, and Pypolca. As shown in **Fig. 1B**, ProGenFixer achieved high sensitivity (97.51%) and precision (99.52%), comparable to the top-performing tools. Furthermore, ProGenFixer demonstrated significant speed advantages, correcting the *E. coli* genome with 100X reads in approximately 12.93 seconds, whereas the fastest competitor, Jasper, required 71.93 seconds. This represents a greater than five-fold speed increase over the other tested tools (**Fig. 1C**). Similar performance gains were observed in simulations using high-coverage (400X) reads (**Fig. 1C**).

Next, we evaluated ProGenFixer’s performance specifically on substitutions and indels independently. To assess ProGenFixer’s capability of substitution correction, we simulated 100X coverage NGS reads with varying substitution rates (0.1‰, 0.3‰, 0.5‰, 0.7‰, 0.9‰, and 1‰). These reads were then processed using ProGenFixer and the comparison tools. As detailed in **Table 1**, ProGenFixer exhibited excellent performance across all substitution rates, achieving high sensitivity (97.54%– 98.49%) and precision (99.56%–100.00%). These achievements were comparable to the best-performing tools in this test, NextPolish and Pypolca.

**Table 1.** Performance of ProGenFixer in handling substitutions. NGS reads with a coverage of 100X were simulated using *dwgsim* (version 0.1.15) with the *E. coli* MG1655 genome sequence (LR881938.1). The expected mutation rates were adjusted using the *-r* option.

|  | ProGenFixer | BWA+GATK | Jasper | NextPolish | Pypolca | Mutation rate |
| --- | --- | --- | --- | --- | --- | --- |
| TP | 456 | 370 | 453 | 460 | 456 | 0.1‰ |
| FP | 0 | 3 | 0 | 0 | 0 |  |
| FN | 7 | 93 | 10 | 3 | 7 |  |
| Sensitivity | 98.49% | 79.91% | 98.49% | 99.35% | 98.49% |  |
| Precision | 100% | 99.20% | 100% | 100% | 100% |  |
| TP | 1332 | 1091 | 1331 | 1352 | 1340 | 0.3‰ |
| FP | 4 | 0 | 1 | 0 | 0 |  |
| FN | 31 | 272 | 32 | 11 | 23 |  |
| Sensitivity | 97.73% | 80.04% | 97.65% | 99.19% | 98.31% |  |
| Precision | 99.70% | 100% | 99.92% | 100% | 100% |  |
| TP | 2319 | 1921 | 2311 | 2352 | 2333 | 0.5‰ |
| FP | 2 | 3 | 2 | 0 | 0 |  |
| FN | 43 | 441 | 51 | 10 | 29 |  |
| Sensitivity | 98.18% | 81.33% | 97.84% | 99.58% | 98.77% |  |
| Precision | 99.91% | 99.84% | 99.91% | 100% | 100% |  |
| TP | 3195 | 2607 | 3188 | 3238 | 3210 | 0.7‰ |
| FP | 13 | 2 | 3 | 0 | 0 |  |
| FN | 64 | 652 | 71 | 21 | 49 |  |
| Sensitivity | 98.04% | 79.99% | 97.82% | 99.36% | 98.50% |  |
| Precision | 99.59% | 99.92% | 99.91% | 100% | 100% |  |
| TP | 4090 | 3337 | 4082 | 4174 | 4127 | 0.9‰ |
| FP | 18 | 8 | 5 | 0 | 0 |  |
| FN | 103 | 856 | 111 | 19 | 66 |  |
| Sensitivity | 97.54% | 79.59% | 97.35% | 99.55% | 98.43% |  |
| Precision | 99.56% | 99.76% | 99.88% | 100% | 100% |  |
| TP | 4448 | 3604 | 4442 | 4521 | 4480 | 1‰ |
| FP | 9 | 7 | 2 | 4 | 0 |  |
| FN | 108 | 952 | 114 | 35 | 76 |  |
| Sensitivity | 97.63% | 79.10% | 97.50% | 99.23% | 98.33% |  |
| Precision | 99.80% | 99.81% | 99.95% | 99.91% | 100% |  |

Note: TP = True Positives, FP = False Positives, FN = False Negatives

We also systematically evaluated ProGenFixer’s performance in correcting indels of varying expected lengths (ranging from~1.1 bp to~20 bp), comparing it against other tools (**Table 2**). The results highlight ProGenFixer’s robust accuracy across different indel sizes. Notably, ProGenFixer maintained consistently high sensitivity (96.52%–99.34%) and precision (99.33%–100.00%) across all tested lengths. In contrast, the performance of some other tools degraded significantly as indel length increased. Jasper, in particular, showed a dramatic drop in sensitivity for longer indels (from 86.94% down to 6.72%). While NextPolish and Pypolca performed well on shorter indels, their sensitivity also decreased for the longest indels tested (~10 bp and~20 bp). BWA+GATK consistently exhibited low sensitivity across all lengths. Crucially, ProGenFixer retained high sensitivity (96.52%) and precision (99.33%) even for the longest simulated indels (~20 bp), demonstrating a key advantage in handling challenging indel errors compared to other tools.

**Table 2.** Performance of ProGenFixer in handling indels of varying lengths. NGS reads were simulated using *dwgsim* (version 0.1.15) in a coverage of 100X using the *E. coli* MG1655 genome sequence (LR881938.1). The expected indel lengths were varied using the *-X* option, which controls the probability of indel extension. The calculation method for determining the expected indel lengths associated with different -X values is detailed in the **Materials and Methods** section.

|  | ProGenFixer | BWA+GATK | Jasper | NextPolish | Pypolca | Indel length |
| --- | --- | --- | --- | --- | --- | --- |
| TP | 1210 | 225 | 1078 | 1232 | 1221 | ~1.11 bp<br>*-X 0.1* |
| FP | 2 | 2 | 1 | 0 | 0 |  |
| FN | 30 | 1015 | 162 | 8 | 19 |  |
| Sensitivity | 97.58% | 18.15% | 86.94% | 99.35% | 98.47% |  |
| Precision | 99.83% | 99.12% | 99.91% | 100.00% | 100.00% |  |
| TP | 1290 | 291 | 909 | 1314 | 1301 | ~1.43 bp<br>*-X 0.3* |
| FP | 3 | 5 | 2 | 2 | 0 |  |
| FN | 35 | 1034 | 416 | 11 | 24 |  |
| Sensitivity | 97.36% | 21.96% | 68.60% | 99.17% | 98.19% |  |
| Precision | 99.77% | 98.31% | 99.78% | 99.85% | 100.00% |  |
| TP | 1325 | 285 | 679 | 1354 | 1339 | ~2 bp<br>*-X 0.5* |
| FP | 2 | 1 | 2 | 0 | 0 |  |
| FN | 38 | 1078 | 684 | 9 | 24 |  |
| Sensitivity | 99.34% | 20.91% | 49.82% | 99.34% | 98.24% |  |
| Precision | 100.00% | 99.65% | 99.71% | 100.00% | 100.00% |  |
| TP | 1286 | 287 | 405 | 1306 | 1298 | ~3.33 bp<br>*-X 0.7* |
| FP | 1 | 5 | 6 | 0 | 0 |  |
| FN | 28 | 1027 | 909 | 8 | 16 |  |
| Sensitivity | 97.87% | 21.84% | 30.82% | 99.39% | 98.78% |  |
| Precision | 99.92% | 98.29% | 98.54% | 100.00% | 100.00% |  |
| TP | 1281 | 275 | 140 | 1304 | 1224 | ~10 bp<br>*-X 0.9* |
| FP | 5 | 5 | 4 | 0 | 34 |  |
| FN | 38 | 1044 | 1179 | 15 | 95 |  |
| Sensitivity | 97.12% | 20.85% | 10.61% | 98.86% | 92.80% |  |
| Precision | 99.61% | 98.21% | 97.22% | 100.00% | 97.30% |  |
| TP | 1193 | 259 | 83 | 1135 | 1050 | ~20 bp<br>*-X 0.95* |
| FP | 8 | 6 | 1 | 32 | 30 |  |
| FN | 43 | 977 | 1153 | 101 | 186 |  |
| Sensitivity | **96.52%** | 20.95% | 6.72% | 91.83% | 84.95% |  |
| Precision | **99.33%** | 97.74% | 98.81% | 97.26% | 97.22% |  |

Note: TP = True Positives, FP = False Positives, FN = False Negatives

### Establishment of a web-based platform for ProGenFixer

To enhance the usability and demonstrate the capabilities of ProGenFixer, we have developed a web-based platform that provides an easy-to-use interface for users to correct genome sequence by uploading NGS data and reference genome sequence. The platform automatically processes the provided reference genome and NGS files and corrects the genome sequence utilizing the ProGenFixer. Upon completion of the correction, the server returns the corrected genome sequence in FASTA format, along with a detailed report of the corrected variations in Variant Call Format (VCF). This user-friendly web server can be accessed at: https://progenfixer.biodesign.ac.cn/.

## Conclusion

ProGenFixer offers a fast, accurate, and user-friendly solution for prokaryotic genome sequence correction. Our benchmarking demonstrates that it operates over five times faster than leading tools such as Jasper, NextPolish, and Pypolca, while maintaining comparable sensitivity and precision for error detection and correction. Notably, ProGenFixer excels in indel correction, showing superior accuracy for long indels (>10 bp). These attributes establish ProGenFixer as a valuable addition to the genomics toolkit, especially for research involving prokaryotic organisms.

While ProGenFixer exhibits high overall accuracy, its performance is currently less optimal in repetitive genomic regions, leading to a noticeable number of false positives and false negatives. Future development could mitigate this limitation by integrating a local alignment-based strategy specifically designed to resolve these complex sequences, thereby improving accuracy without sacrificing the tool’s core computational efficiency.

Furthermore, the high efficiency of ProGenFixer is particularly relevant in the context of modern synthetic biology, where biofoundry platforms, big data analytics, and artificial intelligence are driving rapid advancements in microbial strain optimization [18–20]. This evolving paradigm generates vast numbers of mutant strains, demanding substantial computational resources for genomic analysis. ProGenFixer’s efficient framework directly addresses this challenge, enabling the rapid and precise analysis of large-scale genomic datasets. Consequently, this capability can potentially accelerate the design-build-test-learn cycles of biological engineering, fostering more efficient and innovative progress within data-driven synthetic biology and biofoundry initiatives.

## Materials and Methods

### Simulation of variations, NGS reads and calculation of expected length of indels

All variations and associated NGS reads were simulated with *dwgsim* (v0.1.15) with a range of parameters, allowing for the simulation of different rates of substitutions and indels. The simulation of indels were performed using the *-X* option, which stands for a probability (*p*) of an indel is expected to be extended. When an indel occurs, *dwgsim* uses a certain probability *p* to determine whether the indel will continue to extend during simulation. The probability of the indel stopping at any moment is 1−*p*. After each extension, there is still a chance for further extension, making the problem recursive in nature. The average length of an indel can be calculated using a recursive model based on the probability of extension. The length of the indel can be described as a geometric random variable because we are modeling the number of successes (extensions) before the first failure (when the indel stops extending). Let *E*_*L*_ represent the expected length of an indel, the recursive relationship for the expected length *E*_*L*_ is as follows:

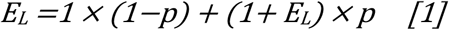

Where: 1×(1−*p*) represents the case where the indel stops after extending once; (1 + *E*_*L*_) × *p* represents the case where the indel continues to extend by one unit and the length is then increased by 1 plus the expected length of any subsequent extensions. By simplifying equation [1], we obtain:

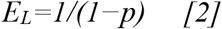

This formula calculates the average indel length, where *p* is the probability of extending the indel by one unit.

### Evaluation Framework

To assess the accuracy of genome correction tools, we developed an evaluation pipeline using Minimap2, SAMTools and BCFtools. The pipeline consists of a bash script (*compare_genomes*.*sh*) and a python script (*vcf_comparison*.*py*). The two scripts are provided alongside the source code of ProGenFixer. The bash script implements a comprehensive workflow for identifying genomic variations between two genome sequences. The process begins with alignment of the query genome to the reference using Minimap2 [21]. The resulting alignments are converted to sorted BAM format using SAMtools, then processed with BCFtools [22] to identify variants, which are exported to VCF format. To evaluate correction accuracy while accounting for the inherent inconsistency in variant representation across different tools, we implemented the following protocol with three key steps: **Step 1: Ground Truth Establishment**: The simulated mutated genome is compared against the original reference genome using compare_genomes.sh to generate a ground truth variant file (VCF file A); **Step 2: Tool Output Analysis**: The corrected genome produced by the correction tool is compared against the same original reference using compare_genomes.sh to generate a prediction variant file (VCF file B); **Step 3: Performance Assessment**: The vcf_comparison.py utility compares a ground truth variant file (VCF file A) against a prediction variant file (VCF file B) using BCFtools isec to calculate key performance metrics. These include: True Positives (TP), representing variants correctly identified; False Positives (FP), representing variants incorrectly identified; False Negatives (FN), representing missed variants; Sensitivity, the proportion of actual variants correctly identified (TP/(TP+FN)); and Precision, the proportion of identified variants that are correct (TP/(TP+FP)).

## Author Contributions

The project’s conceptualization and experimental design were carried out by LFS, with overall supervision provided by JSL, PZ, JBS and YHM. LFS devised the algorithms and conducted the computational analyses. MW, XMN, LLL and BTH participated in the simulation analysis. XPL, ZKS, RYW, HRL and HWM developed the web platform. The manuscript was written by LFS, with valuable contributions and insights provided by all co-authors.

## Acknowledgments

This research was supported by the Tianjin Synthetic Biotechnology Innovation Capacity Improvement Project (TSBICIP-CXRC-072) and (TSBICIP-PTJJ-012).

## Data and Code Availability

ProGenFixer is freely available at https://github.com/Scilence2022/ProGenFixer.

